# Cross-cancer knowledge transfer for lymph node metastasis prediction: Applicability of an axillary lymph node-based deep learning model

**DOI:** 10.64898/2026.11.04.764191

**Authors:** Jaewoong Kang, Jeong Wook Kang, Lyo Min Kwon, Bum-Joo Cho, Bum Jun Kim

## Abstract

Evaluation of lymph node (LN) metastasis is critical for cancer staging and treatment planning. Since radiologic hallmarks of metastatic LNs are shared across primary cancers, a computed tomography (CT)-based deep learning (DL) classifier developed from one anatomy may transfer to others. This study built a DenseNet-121 ensemble model using clinical-implication-applied preprocessing and evaluated its cross-anatomical generalizability to LNs from diverse primary cancers. A DenseNet-121 ensemble model combining fixed-size cropping (FSC) and adjustable square cropping (ASC) was trained from ImageNet initialization on an axillary lymph node (ALN) dataset (n = 1,128 from 523 breast cancer patients). The model utilized clinical-implication-applied perinodal cropping with bicubic filtering and standardized resizing. To test cross-anatomical generalizability, an external dataset of 256 CT images across four anatomical regions was evaluated: cervical, mediastinal, upper-abdominal, and lower-abdominal LNs. Perinodal free margins were systematically varied from 10% to 40% to optimize the balance between context preservation and surround interference. Bicubic filtering and standardized resizing yielded the highest internal performance on the original ALN dataset (FSC-ASC ensemble area under the receiver operating characteristic curve [AUROC] 0.958, 95% confidence interval [CI] 0.920–0.987 at 40% margin). The model demonstrated site-dependent, above-chance generalizability across all external regions. Cervical LNs achieved the highest performance at a 40% margin (AUROC 0.939, 95% CI 0.875–0.989). Mediastinal LNs peaked at a 10% margin (AUROC 0.796, 95% CI 0.672–0.903), with wider margins degrading performance due to adjacent structural interference. Upper-abdominal and lower-abdominal LNs reached optimal performance at a 30% margin (AUROC 0.814 for upper; AUROC 0.917 for lower). A single ALN-trained DL model can accurately predict LN metastasis across cervical, mediastinal, and abdominal stations through site-specific perinodal margin tuning. These findings support the feasibility of cross-cancer knowledge transfer and the clinical utility of generalizable backbones for pan-cancer nodal assessment.

## Introduction

The presence and extent of lymph node (LN) metastasis are critical determinants for clinical staging, prognostic evaluation, and treatment planning across most solid cancers [1,2].

Traditionally, diagnostic criteria for metastatic LNs in clinical practice have primarily relied on imaging-based morphological features, with size—specifically a short-axis diameter exceeding 1 cm—serving as the most widely accepted threshold [3,4]. However, this approach is limited by interobserver variability and low sensitivity. A significant proportion of metastatic LNs maintain a normal size, which can lead to the clinical underestimation of cancer stages and subsequent mistreatment [5–7].

A “clinical-implication-applied” perinodal cropping strategy for computed tomography (CT)-based axillary lymph node (ALN) classification has previously been shown in our previous study to outperform conventional size-based criteria [8]. However, whether the malignancy signal captured by the ALN-based classifier is transferable to LNs of other anatomical regions from diverse cancers has not been systematically tested. Because metastatic LNs across different anatomical regions share radiologic hallmarks such as cortical thickening and loss of the fatty hilum, the same convolutional features may carry over to other anatomical stations, providing a basis for cross-cancer knowledge transfer.

While metastatic LNs across different anatomical regions share certain radiologic hallmarks, such as cortical thickening or loss of the fatty hilum [9,10], their surrounding environments vary significantly. For instance, cervical, mediastinal, and abdominal LNs are encased by complex anatomical structures—including major airways, great vessels, and digestive organs—which may interfere with precise feature extraction when compared to the relatively isolated and fatty environment of the axilla [11–13]. Given these anatomical differences, whether an ALN-based deep learning (DL) model can be generalized to other anatomical sites—i.e., whether cross-cancer knowledge transfer is feasible—remains unclear.

In this study, we first verified the predictive performance of a DenseNet-121-based DL model— sharpened with bicubic interpolation and standardized resizing—on the ALN dataset. We then assessed its applicability to four independent external cohorts comprising cervical, mediastinal, upper-abdominal, and lower-abdominal LNs from patients with head and neck, lung, gastric, and colorectal cancers, respectively. By treating the perinodal margin as a tunable hyperparameter and comparing site-specific operating points, we sought to delineate the anatomical factors that govern the cross-cancer applicability of an ALN-based DL model, demonstrating that site-specific perinodal margin tuning enables accurate cross-anatomical metastasis prediction.

## Materials and methods

### Study population

The development cohort was the curated ALN dataset previously established for breast cancer ALN classification [8]; reusing it ensured comparability with the published benchmark. To assess cross-cancer applicability, we then assembled four prospectively independent external validation cohorts spanning four different primary cancers and anatomical stations. The four external cohorts comprised consecutive patients who underwent preoperative contrast-enhanced CT at Hallym University Medical Center between January 2012 and July 2022 and were eligible if they had (i) a histologically confirmed primary tumor of the head and neck, lung, stomach, or colorectum, (ii) a high-resolution contrast-enhanced preoperative CT scan, and (iii) a clear anatomical correspondence between the radiologic target LN and its pathology report. The benign or malignant status of every LN was confirmed pathologically from a surgical specimen or a percutaneous core-needle biopsy. Patients with non-enhanced or low-dose CT scans, or with prior neoadjuvant chemotherapy or radiotherapy, were excluded to avoid imaging-quality and treatment-induced confounds. The study was reviewed and approved by the Institutional Review Board of Hallym University Medical Center (registration number: HALLYM 2023-07-017; date of approval: 5 September 2023) in accordance with the Declaration of Helsinki, and the requirement for written informed consent was waived by the Institutional Review Board because the analysis used fully anonymized retrospective data.

### Image acquisition and reconstruction

Imaging was performed on 64- to 128-row multidetector CT systems (scanners from Siemens Healthineers and Philips Healthcare). Scanning parameters included peak voltages between 80 and 140 kVp, tube currents spanning 45–714 mAs, pitch values of either 0.6 or 1.2, and collimations of 128 × 0.6 mm or 192 × 0.6 mm. Axial images were reconstructed at a 3-mm slice thickness with pixel dimensions of 0.55–0.99 mm using the Advanced Modeled Iterative Reconstruction (ADMIRE-v2) algorithm (Siemens Healthineers).

### Image preprocessing strategies

The LN image preprocessing protocol was strategically designed to capture alterations in the perinodal soft tissue. To incorporate the perinodal context, the bounding box was generated to include a free margin of 40% of the LN’s maximum diameter (Fig 1). Within this protocol, two complementary cropping methods were used—the Fixed Size Crop (FSC) method, which preserves the LN’s actual physical dimensions to retain size-dependent features, and the Adjustable Square Crop (ASC) method, which standardizes the input to a fixed pixel grid to mitigate size bias and emphasize internal texture and subtle perinodal changes. The two methods were combined by averaging their predicted malignancy probabilities to form the FSC-ASC ensemble model.

**Fig 1.**
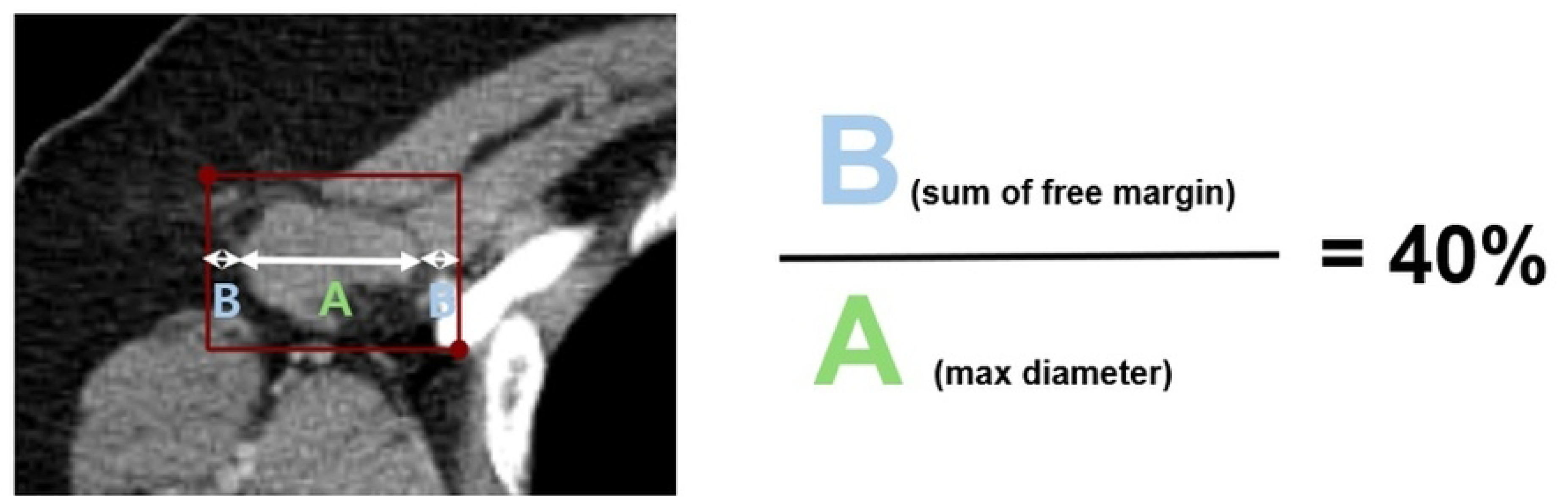
Preprocessing protocol for clinical implication-applied lymph node segmentation. The schematic illustrates the generation of a bounding box incorporating a free margin corresponding to exactly 40% of the target lymph node’s maximum diameter, establishing a unified standard across physical and pixel coordinates.

### Development of the optimized ALN-based DL model

To enhance the baseline ALN prediction framework and develop an optimized model, two primary image-refinement strategies were implemented (Fig 2). First, bicubic interpolation was adopted over standard bilinear schemes to enhance the perceptual clarity of LN morphology and texture. This filtering method mitigates aliasing artifacts and produces smoother intensity transitions, thereby preserving critical structural details more effectively than linear methods [14]. Second, cropped regions of interest were resized to a fixed target resolution to address the problem that upsampling can increase low-information background regions. This standardized resizing step minimizes computational overhead and stabilizes batch training by ensuring consistent feature representation across different scales [15,16].

**Fig 2.**
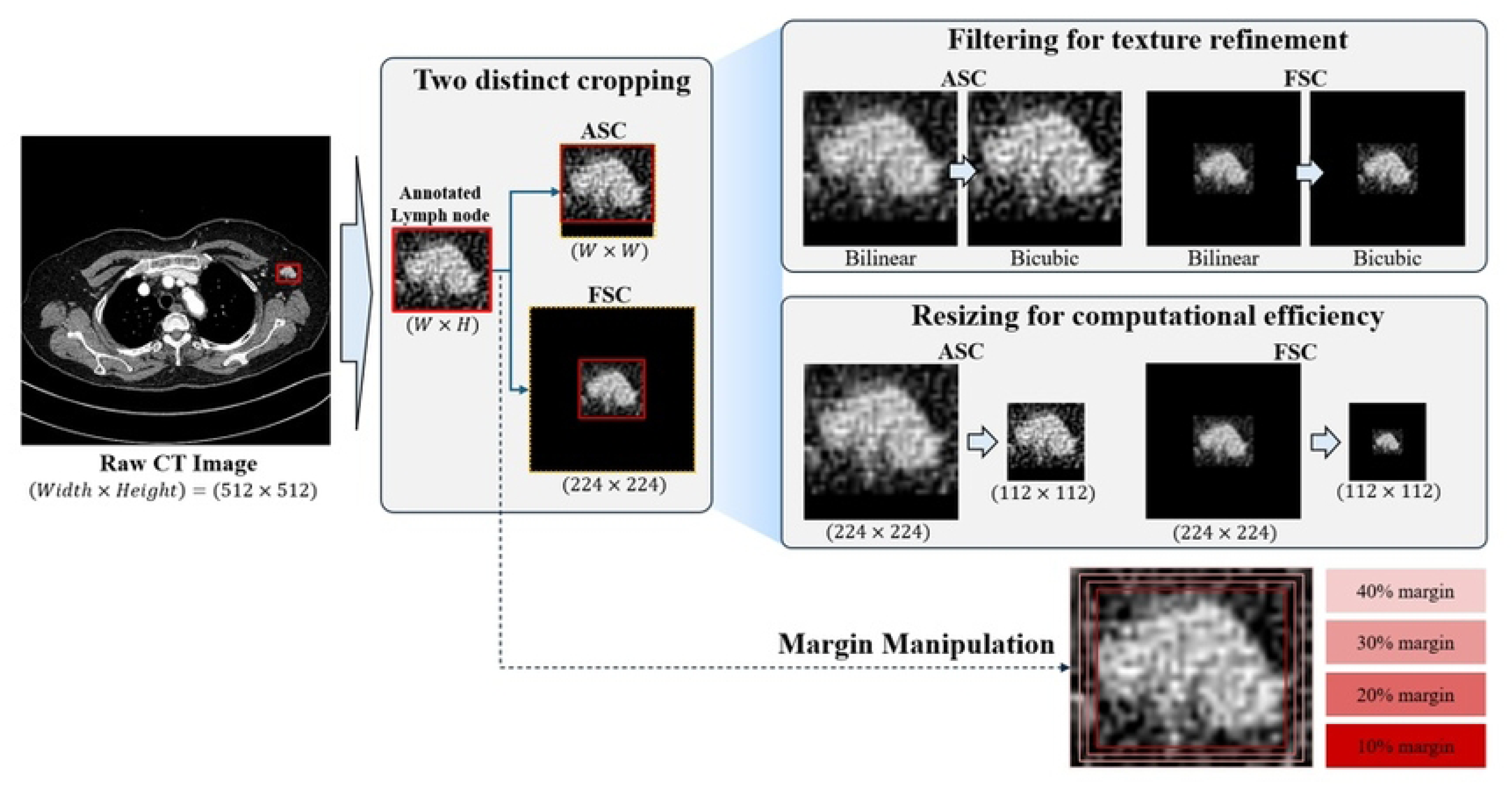
Overview of the optimized axillary lymph node-based deep learning framework. The image reconstruction pipeline processes raw CT input (512 × 512 matrix) through two cropping layers: Fixed Size Crop (FSC) to retain absolute structural proportions, and Adjustable Square Crop (ASC) to normalize internal patterns. Texture resolution is enhanced via bicubic filtering and standardized downsampling matrix grids before prediction averaging inside the DenseNet-121 ensemble network layer.

The optimization process involved evaluating these strategies independently within both ASC and FSC frameworks. We selected the specific configurations that yielded the highest diagnostic performance for each cropping method. These top-performing components were then integrated using an ensemble method to construct the final optimized model. To validate the efficacy of this refinement, the performance of the ensemble model was benchmarked against the published results [8] on the same ALN dataset.

### Cross-anatomical performance evaluation

The optimized model’s ability to differentiate between malignant and benign LNs across different anatomical sites was evaluated using four independent groups: cervical LNs in head and neck cancer, mediastinal LNs in lung cancer, upper-abdominal LNs in gastric cancer, and lower-abdominal LNs in colorectal cancer. Performance at each site was reported separately for the FSC method, the ASC method, and the FSC-ASC ensemble.

A critical challenge in cross-anatomical application is the interference from complex anatomical structures, such as major airways, great vessels, and digestive organs, which vary significantly by region. Whereas the ALN-based DL model was developed at a fixed 40% margin, here we treated the perinodal margin as a tunable hyperparameter and varied the free margin from 10% to 40%. While a larger margin captures broader contextual information, it risks diluting the target signal or introducing noise from adjacent non-lymphatic structures. The optimum margin per anatomical site was therefore determined by grid search, balancing contextual coverage and target fidelity—a design choice consistent with recent findings that multi-scale spatial context is critical for small-lesion analysis in volumetric data [17,18].

### Network architecture and training

We employed a DenseNet-121 backbone, which was previously shown to be effective for malignant LN discrimination on CT [8]. DenseNet-121 was selected for its efficient feature reuse and improved gradient flow, which have proven effective in medical image classification tasks [19].

Although the task is a binary classification problem, categorical cross-entropy loss was selected as the loss function because categorical cross-entropy can enhance sensitivity for detecting abnormalities compared to standard binary loss functions [20]. The network was optimized using the Adam optimizer, a widely adopted algorithm in computer vision, with an initial learning rate of 1 × 10−4. A learning rate scheduler was applied to decay the rate by a factor of 0.1 every 10 epochs.

To prevent overfitting, early stopping was implemented with a patience of 10 epochs, triggered after the first 20 epochs. The training process was capped at 100 epochs, and the model checkpoint achieving the lowest validation loss was retained as the final model. Model training was performed on a workstation distinct from the original ALN-based DL model [8] setup, equipped with an Intel Xeon Silver-class central processing unit and an NVIDIA GeForce RTX 4090 graphics processing unit under a custom liquid-cooling enclosure. The change in compute environment, together with an updated training dataset (n = 1,128 vs. n = 1,127 in the original ALN-based DL model [8]), means that all weights reported here were trained from ImageNet initialization rather than fine-tuned from previously released checkpoints.

### Statistical analyses

For both the internal ALN test split and the external multi-site cohort, predictive performance was summarized by accuracy, area under the receiver operating characteristic curve (AUROC), sensitivity, specificity, positive predictive value, negative predictive value, and the F1 score.

Operating points were chosen by maximizing the Youden J statistic on the validation split. Two-sided 95% confidence intervals (CIs) for every metric were obtained by case-resampled bootstrap with 1,000 replicates, drawn at the LN level. Differences in AUROC across the four enhancement strategies (Baseline, Filtering, Resizing, and Filtering + Resizing) within each crop family were compared by one-way analysis of variance (ANOVA), separately for the FSC and ASC methods. Results from the FSC and ASC methods were integrated by averaging their predicted malignancy probabilities to form the FSC-ASC ensemble. Cross-anatomical predictions were generated by applying this ensemble, without any site-specific re-training, to each external site at perinodal margins of 10%, 20%, 30%, and 40%; the per-site optimum margin was the one that maximized the bootstrap-mean AUROC. A two-sided p < 0.05 was considered statistically significant. All statistical analyses were performed in Python (version 3.10) using scikit-learn 1.5, SciPy 1.14, and NumPy 1.26.

## Results

### Performance of original model on the ALN dataset

We first re-evaluated the performance of the DenseNet-121 ensemble on the held-out ALN test split (n = 112) before applying it to other anatomical sites, because the present model was trained on a slightly expanded ALN dataset (n = 1,128) [8]. Across the four perinodal margins (10%, 20%, 30%, and 40%) and the three prediction methods (FSC, ASC, FSC-ASC ensemble), AUROC ranged from 0.897 to 0.960 with bootstrap (1,000 iterations) 95% CIs (S1 Table).

Performance increased monotonically with the perinodal margin for the ensemble method (AUROC 0.920 at 10%, 0.925 at 20%, 0.956 at 30%, and 0.958 at 40%) and peaked at the 40% margin for both the FSC method (AUROC 0.960, 95% CI 0.920–0.990) and the ensemble (AUROC 0.958, 95% CI 0.920–0.987). The ensemble’s 95% CI at the 40% margin (0.920– 0.987) overlapped the previously published estimate (AUROC 0.968) reported by Park et al. [8], supporting that the re-trained model reproduces the original performance benchmark and is therefore a valid backbone for the cross-anatomical experiments that follow.

### Optimization of model: Effect of bicubic interpolation and standardized resizing

To establish the optimal preprocessing framework, the optimized model was first validated using the original ALN dataset (n = 1,128). The integration of bicubic filtering and standardized resizing yielded the highest diagnostic performance across both cropping strategies (Table 1).

**Table 1.** Diagnostic performance of the optimized model according to two different strategies (bicubic filtering and standardized resizing) for Fixed Size Crop (FSC) and Adjustable Square Crop (ASC) methods.

| **Crop** | **Method** | **Accuracy** | **AUROC** | **Recall** | **Precision** | **p-value\*** |
|---|---|---|---|---|---|---|
| FSC | Baseline | 0.889 ± 0.022 | 0.950 ± 0.013 | 0.884 ± 0.059 | 0.915 ± 0.043 | 0.90 |
| FSC | Filtering alone | 0.897 ± 0.024 | 0.953 ± 0.013 | 0.904 ± 0.055 | 0.911 ± 0.037 | 0.90 |
| FSC | Resizing alone | 0.895 ± 0.015 | 0.953 ± 0.011 | 0.874 ± 0.038 | 0.933 ± 0.032 | 0.90 |
| FSC | Filtering + Resizing | 0.896 ± 0.024 | 0.954 ± 0.013 | 0.852 ± 0.050 | 0.954 ± 0.024 | 0.90 |
| ASC | Baseline | 0.913 ± 0.024 | 0.954 ± 0.016 | 0.893 ± 0.041 | 0.947 ± 0.029 | 0.63 |
| ASC | Filtering alone | 0.912 ± 0.022 | 0.956 ± 0.015 | 0.889 ± 0.050 | 0.949 ± 0.025 | 0.63 |
| ASC | Resizing alone | 0.907 ± 0.026 | 0.949 ± 0.021 | 0.884 ± 0.038 | 0.943 ± 0.023 | 0.63 |
| ASC | Filtering + Resizing | 0.917 ± 0.024 | 0.959 ± 0.017 | 0.903 ± 0.043 | 0.944 ± 0.031 | 0.63 |

Data are presented as mean ± standard deviation. \*AUROC values of the four preprocessing configurations within each crop method were compared using one-way analysis of variance (ANOVA). ASC, adjustable square crop; AUROC, area under the receiver operating characteristic curve; FSC, fixed-size crop.

For the FSC method, the combined strategy achieved an AUROC of 0.954 ± 0.013 and an accuracy of 0.896 ± 0.024, showing a numerical improvement over the baseline AUROC of 0.950 ± 0.013. For the ASC method, the combined strategy reached an AUROC of 0.959 ± 0.017 and an accuracy of 0.917 ± 0.024, surpassing the baseline AUROC of 0.954 ± 0.016. Statistical analysis using one-way ANOVA indicated no significant difference in AUROC among the methods (*p* = 0.90 for FSC; *p* = 0.63 for ASC). These results suggest that while the baseline model was already highly robust, the augmented strategies further stabilized and refined the predictive performance for cross-cancer knowledge transfer.

### External validation cohort: Patient and image characteristics

To evaluate the model’s generalizability, a total of 256 LN images were collected from 98 patients (24, 24, 27, and 23 patients with head and neck, lung, gastric, and colorectal cancer, respectively) (Table 2). The dataset was distributed across four distinct anatomical regions and primary malignancies: head and neck cancer (cervical LNs, n = 67), lung cancer (mediastinal LNs, n = 64), gastric cancer (upper-abdominal LNs, n = 61), and colorectal cancer (lower-abdominal LNs, n = 64). The malignancy rate was maintained at 50% across all regions, with 128 malignant and 128 benign LNs pathologically confirmed by core-needle biopsy or surgery.

**Table 2.** Characteristics and distribution of the multi-site external lymph node dataset.

| **Metric** | **Cervical LNs** | **Mediastinal LNs** | **Upper-abdominal LNs** | **Lower-abdominal LNs** |
|---|---|---|---|---|
| No. of Patients | 24 (head and neck cancer) | 24 (lung cancer) | 27 (gastric cancer) | 23 (colorectal cancer) |
| No. of Images (Overall) | 67 | 64 | 61 | 64 |
| Malignant | 32 | 33 | 31 | 32 |
| Benign | 35 | 31 | 30 | 32 |

LN, lymph node.

### Cross-regional predictive performance of optimized model

The FSC-ASC ensemble was then applied to evaluate diagnostic accuracy across the four external anatomical sites (Table 3 and S2 Table). Fig 3 provides the corresponding receiver operating characteristic (ROC) curves stratified by perinodal margin. Performance was strongly modulated by the perinodal margin size, reflecting the impact of regional anatomical complexity.

**Fig 3.**
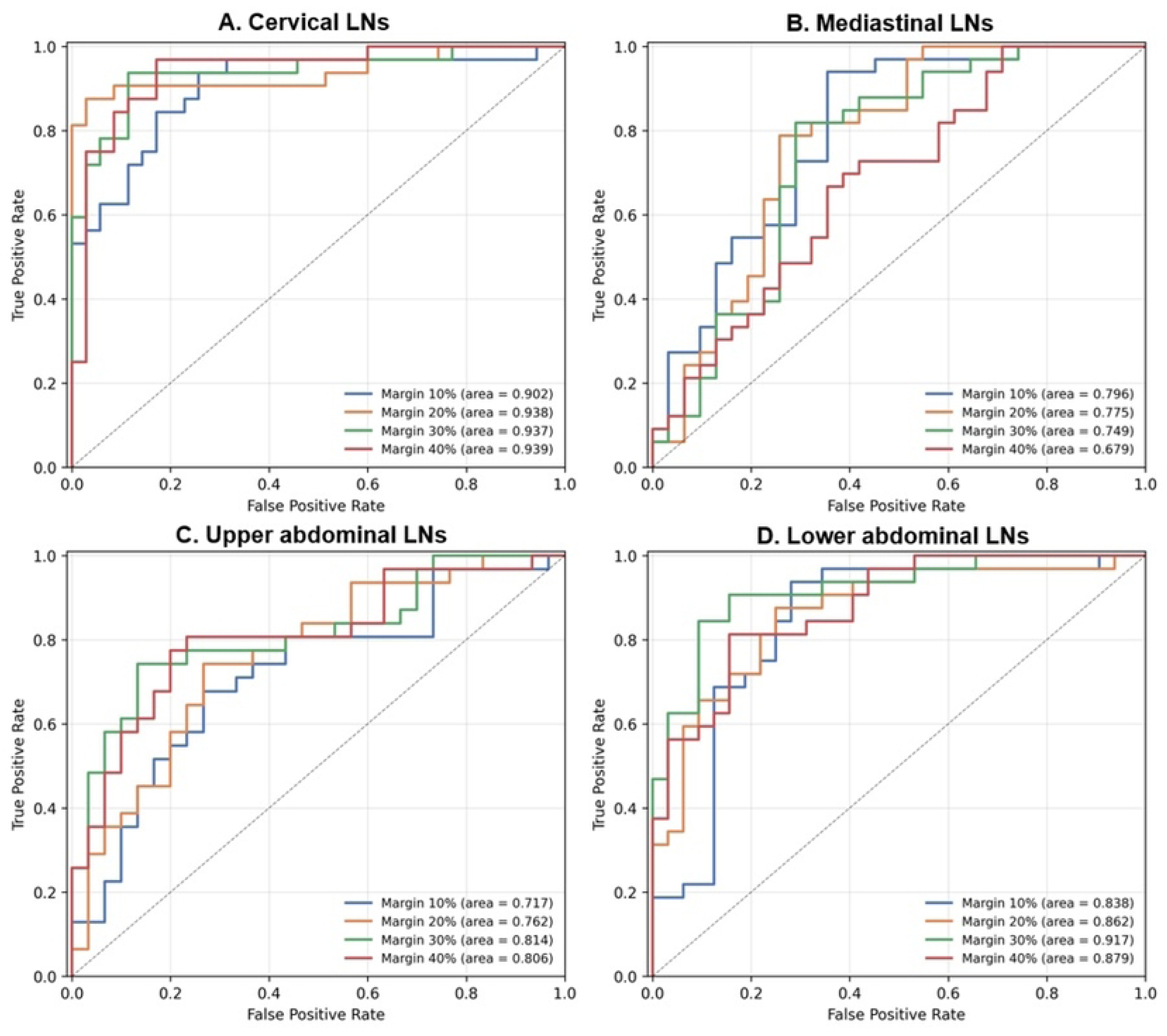
Receiver operating characteristic (ROC) analysis across independent external validation sites. ROC curves illustrate diagnostic performance across regional cohorts stratified by perinodal free margins (10%, 20%, 30%, and 40%): (A) Cervical lymph nodes, (B) Mediastinal lymph nodes, (C) Upper-abdominal lymph nodes, and (D) Lower-abdominal lymph nodes. True positive rates are plotted against false positive rates to highlight the distinct site-dependent impact of peripheral contextual structures.

**Table 3.** Predictive performance metrics of the optimized ALN-based deep learning ensemble model across external validation cohorts stratified by perinodal free margin.

| Region<br>(Cancer Type) | Metric | Margin 40% | Margin 30% | Margin 20% | Margin 10% |
|---|---|---|---|---|---|
| **Cervical LNs (head and neck cancer)** | AUROC | **0.939 ± 0.029** | 0.937 ± 0.030 | 0.938 ± 0.033 | 0.902 ± 0.037 |
| **Cervical LNs (head and neck cancer)** | Accuracy | 0.895 ± 0.034 | 0.910 ± 0.034 | 0.925 ± 0.030 | 0.836 ± 0.042 |
| **Mediastinal LNs (lung cancer)** | AUROC | 0.679 ± 0.065 | 0.749 ± 0.065 | 0.775 ± 0.061 | **0.796 ± 0.057** |
| **Mediastinal LNs (lung cancer)** | Accuracy | 0.656 ± 0.056 | 0.766 ± 0.052 | 0.766 ± 0.052 | 0.797 ± 0.052 |
| **Upper-abdominal LNs (gastric cancer)** | AUROC | 0.806 ± 0.057 | **0.814 ± 0.056** | 0.762 ± 0.060 | 0.717 ± 0.067 |
| **Upper-abdominal LNs (gastric cancer)** | Accuracy | 0.787 ± 0.050 | 0.803 ± 0.050 | 0.738 ± 0.054 | 0.705 ± 0.059 |
| **Lower-abdominal LNs (colorectal cancer)** | AUROC | 0.879 ± 0.041 | **0.917 ± 0.035** | 0.862 ± 0.045 | 0.838 ± 0.054 |
| **Lower-abdominal LNs (colorectal cancer)** | Accuracy | 0.828 ± 0.048 | 0.875 ± 0.044 | 0.812 ± 0.048 | 0.828 ± 0.048 |

Data are presented as bootstrap mean ± standard deviation. Bolded values indicate the optimum operating point determined by grid search for each anatomical region. ALN, axillary lymph node; AUROC, area under the receiver operating characteristic curve; LN, lymph node.

Cervical LNs demonstrated the highest regional generalizability, with a peak AUROC of 0.939 at a 40% margin, suggesting that broader contextual information is beneficial in the relatively accessible soft tissue of the neck. In contrast, mediastinal LNs showed the most distinct trend, with performance increasing as the margin size decreased. The ensemble reached an AUROC of 0.796 (95% CI 0.672–0.903) at the 10% margin, whereas the 40% margin yielded the lowest performance (AUROC 0.679, 95% CI 0.537–0.805). For the FSC method the gap was even larger, and was the only across-margin comparison in the cross-anatomical cohort to reach this level of significance. This is consistent with interference from adjacent airways and major vessels degrading the model when the perinodal margin is too wide. Both upper-and lower-abdominal LNs peaked at the 30% margin (AUROC 0.814, 95% CI 0.684–0.912 for upper; AUROC 0.917, 95% CI 0.849–0.977 for lower), indicating a balance between nodal features and contextual fat planes.

Per-LN error counting across the 12 prediction methods (4 margins × {FSC, ASC, FSC-ASC ensemble}) revealed that hard cases—LNs misclassified by at least 8 of the 12 methods— accounted for 4% of cervical LNs, 20% of mediastinal LNs, 20% of upper-abdominal LNs, and 13% of lower-abdominal LNs.

### Per-site classification performance

At the per-site optimum perinodal margin, the ensemble achieved AUROC = 0.939 ± 0.029 (cervical, 40% margin, n = 67), 0.796 ± 0.057 (mediastinal, 10% margin, n = 64), 0.814 ± 0.056 (upper-abdominal, 30% margin, n = 61), and 0.917 ± 0.035 (lower-abdominal, 30% margin, n = 64), with corresponding accuracies of 0.896 ± 0.034, 0.797 ± 0.052, 0.803 ± 0.050, and 0.875 ± 0.045 (S2 Table). At the Youden J operating threshold, sensitivities ranged from 0.742 ± 0.079 (upper-abdominal) to 0.969 ± 0.026 (cervical) and specificities from 0.645 ± 0.081 (mediastinal) to 0.906 ± 0.054 (lower-abdominal); the corresponding optimal probability thresholds spanned 0.587 (mediastinal) to 0.874 (upper-abdominal), confirming that a single decision threshold transferred from the ALN would not be appropriate for cross-cancer use.

## Discussion

This study tested whether a CT-based DenseNet-121 ensemble, sharpened with bicubic interpolation and standardized resizing, can be applied to LNs drained from non-axillary anatomical regions from diverse primary cancers. The ensemble achieved site-dependent but consistently above-chance performance across cervical, mediastinal, and abdominal LNs, and the optimum perinodal margin differed between anatomical stations. Together these findings establish per-site margin tuning as a decisive variable in the cross-cancer applicability of CT-based LN classifiers.

Traditionally, the radiologic diagnosis of LN metastasis relies on common malignant signs regardless of the anatomical site, such as cortical thickening, loss of the fatty hilum, and transition to a rounded shape. The fact that the optimized ALN-based DL model achieved excellent performance across different sites suggests that these radiologic hallmarks serve as universal feature extractors in the DL framework. This goes beyond the concept of “universal lesion detection” proposed by the National Institutes of Health DeepLesion study [21], demonstrating the feasibility of “Cross-cancer Knowledge Transfer” where malignancy-specific knowledge learned from one cancer type can be effectively applied to others [22]. These findings provide a strong rationale for developing efficient multi-site models by leveraging transfer learning from data-rich anatomical stations, rather than training individual models for every specific site from scratch.

A key finding is that the efficiency of this knowledge transfer depends on the balance between essential contextual information and anatomical noise, which varies by site:

### Cervical LNs

The highest performance (AUROC 0.939) was achieved with a wide margin of 40%. Similar to the axillary region, the neck is rich in peripheral fat and soft tissue. Including a wider margin likely allowed the model to better capture subtle irregularities at the LN interface or perinodal infiltration, which Som et al. and Kann et al. have emphasized as critical indicators of metastasis [23,24].

### Mediastinal LNs

This region showed the most dramatic performance shift, where AUROC improved from 0.679 at the 40% margin to 0.796 at the 10% margin. The reversal of the cervical pattern provides the strongest single piece of evidence in this work that the optimal perinodal margin is site-dependent rather than universal, and is consistent with complex structures such as the airway, great vessels, and esophagus acting as interference noise when included in a wide margin, hindering the extraction of LN-specific features [25].

### Abdominal LNs

Both gastric (upper) and colorectal (lower) cancer LNs showed optimal performance at a 30% margin. This reflects the anatomical characteristic of the abdomen, where a balance must be struck between capturing the LN’s internal features and the surrounding mesenteric fat while excluding interference from adjacent visceral organs [26,27].

Our optimized ALN-based DL model demonstrated diagnostic performance that is comparable to, or in some instances exceeds, that of site-specific models reported in the literature. For example, while Zhang et al. reported an AUROC of 0.82–0.87 for a DL model specifically trained on gastric cancer LNs [28], our ensemble achieved a per-site optimum AUROC range of 0.814–0.917 for abdominal LNs, despite its cross-anatomical application. Notably, compared to the recent study by Chu and Fishman, which reported an AUROC of 0.92 for predicting pancreatic LN metastasis using a site-specific artificial intelligence model [29], our ALN-based model reached a per-site optimum AUROC of 0.939 in cervical LNs and 0.917 in lower-abdominal LNs without any site-specific re-training, and was slightly below site-specific performance in mediastinal (0.796) and upper-abdominal (0.814) LNs. Together, these comparisons indicate that a single ALN-based DL model can be leveraged across multiple primaries with site-aware perinodal-margin tuning, which is the operational essence of cross-cancer knowledge transfer for LN metastasis.

Technically, the superior diagnostic performance observed in this study was again driven by the combination of clinical-implication-applied perinodal cropping (FSC plus ASC, integrated through ensemble averaging) and the bicubic filtering and standardized resizing strategies.

Extending the model from the axilla to the four external sites further indicates that these preprocessing choices, rather than anatomy-specific training, are the dominant factors in cross-cancer knowledge transfer of CT-based LN classifiers.

This study has several limitations. First, it is a retrospective single-center analysis whose 256-LN external cohort, although spanning four anatomical regions, remains modest in absolute size; multi-center validation is needed to confirm robustness across different scanner vendors and image-reconstruction pipelines. Second, end-to-end clinical deployment will require coupling the present classifier with an automated LN-localization step, which we did not evaluate here, since all bounding boxes were drawn manually by a single reference radiologist. Third, our analysis was restricted to contrast-enhanced CT; whether the same cross-cancer knowledge-transfer framework generalizes to ultrasound or magnetic resonance imaging—modalities that are routinely used for cervical, breast, and abdominal LN evaluation—is an important direction for future work.

## Conclusions

In conclusion, an ALN-based DL model trained solely on axillary LNs from breast cancer patients transfers to cervical, mediastinal, and abdominal LNs with site-dependent but consistently above-chance performance, supporting the feasibility of cross-cancer knowledge transfer for LN metastasis prediction. Furthermore, the per-site optimum perinodal margin identified here also offers a concrete tuning guide for researchers extending CT-based LN classifiers beyond breast cancer.

## Acknowledgments

None.

## Author contributions

**Conceptualization:** Bum-Joo Cho, Bum Jun Kim.

**Data curation:** Lyo Min Kwon, Bum Jun Kim.

**Formal analysis:** Jaewoong Kang.

**Funding acquisition:** Bum-Joo Cho, Bum Jun Kim.

**Investigation:** Lyo Min Kwon, Bum-Joo Cho, Bum Jun Kim.

**Methodology:** Jaewoong Kang, Bum-Joo Cho.

**Project administration:** Bum Jun Kim.

**Resources:** Jeong Wook Kang, Lyo Min Kwon.

**Software:** Jaewoong Kang.

**Supervision:** Bum-Joo Cho.

**Validation:** Jeong Wook Kang, Bum-Joo Cho, Bum Jun Kim.

**Visualization:** Jaewoong Kang.

**Writing – original draft:** Jaewoong Kang, Jeong Wook Kang.

**Writing – review & editing:** Jaewoong Kang, Lyo Min Kwon, Bum-Joo Cho, Bum Jun Kim.

## Supporting information

**S1 Table. Axillary cohort—performance of the FSC, ASC, and FSC-ASC ensemble methods across four different perinodal margins on the held-out test split (n = 112).** Data are presented as point estimate (95% confidence interval). ASC, adjustable square crop; AUROC, area under the receiver operating characteristic curve; F1, F1 score; FSC, fixed-size crop; NPV, negative predictive value; PPV, positive predictive value; Thresh., optimal probability threshold.

(DOCX)

**S2 Table. Cross-anatomical performance of optimized ALN-based DL model: Full classification metrics across margins.** Data are presented as point estimate (95% confidence interval). ALN, axillary lymph node; AUROC, area under the receiver operating characteristic curve; DL, deep learning; F1, F1 score; LN, lymph node; NPV, negative predictive value; PPV, positive predictive value.

(DOCX)

